# Piezo2 expressing nociceptors mediate mechanical sensitization in experimental osteoarthritis

**DOI:** 10.1101/2022.03.12.484097

**Authors:** Alia M. Obeidat, Matthew J. Wood, Shingo Ishihara, Jun Li, Lai Wang, Dongjun Ren, David A. Bennett, Richard J. Miller, Anne-Marie Malfait, Rachel E. Miller

## Abstract

Osteoarthritis is a very common painful joint disease, for which few treatment options exist. New non-opioid targets are needed for addressing osteoarthritis pain, which is mechanical in nature and associated with daily activities such as walking and climbing stairs. Piezo2 has been implicated in development of mechanical pain, but the mechanisms by which this occurs remain poorly understood. We observed that in two different murine models of osteoarthritis (destabilization of the medial meniscus and natural aging), nociceptor-specific *Piezo2* conditional knock-out mice developed osteoarthritic joint damage, but were protected from associated mechanical sensitization. Since nerve growth factor (NGF) is known to mediate nociceptor sensitization, and antibodies that neutralize NGF are effective as a treatment for osteoarthritis pain, we explored the effects of intra-articularly injected NGF on the development of mechanical joint pain. Wild-type mice developed knee swelling and mechanical pain in response to intra-articular NGF, while nociceptor-specific *Piezo2* conditional knock-out mice were protected from these effects. Single cell RNA sequencing and *in situ* hybridization of mouse and human lumbar dorsal root ganglia (DRG) revealed that a subset of nociceptors co-express *Piezo2* and *Ntrk1* (the gene that encodes the NGF receptor TrkA). These results indicate that Piezo2 plays a key role in nociceptor sensitization processes in the osteoarthritic joint, and targeting Piezo2 may represent a novel therapy for osteoarthritis pain control.

**One Sentence Summary:** Nociceptor sensitization to mechanical stimuli is dependent on Piezo2 in mouse models of osteoarthritis.

## INTRODUCTION

Nociceptors are sensory neurons that enable the body to sense noxious chemical, mechanical and thermal stimuli to avoid tissue damage. Under pathologic conditions, such as during inflammation, nociceptors become sensitized so that they respond to normally non-noxious stimuli. The concept of nociceptor sensitization has been extensively described, but the precise molecular underpinnings of sensitization to mechanical stimuli are incompletely understood. Recent work suggests that the mechanosensitive ion channel, Piezo2, contributes to a range of fundamental sensory neuron functions, including sensing of light touch (*1*), proprioception (*2*), vibratory detection (*3*), and more recently, mechanical pain (*3, 4*). In particular, it was demonstrated that in models of acute inflammation or nerve injury, *Piezo2* knock-out mice showed reduced mechanical allodynia (*4*). Furthermore, individuals with PIEZO2 loss of function mutations were unable to detect light touch applied to skin after inflammation was induced with capsaicin (*3*).

Although it seems increasingly clear that Piezo2 may mediate mechanical sensitization induced by inflammation in mouse models, it is not clear whether this phenomenon contributes to pain associated with specific diseases. Osteoarthritis, one of the world’s most common diseases, is characterized by mechanically driven pain in the presence of tissue injury and low-grade inflammation (*5*). As the most common form of arthritis, osteoarthritis is one of the leading sources of chronic pain affecting 240 million people worldwide (*6-8*). Clinical research has demonstrated that people afflicted with osteoarthritis develop mechanical sensitization not only at the affected joint but also at sites away from the joint, suggesting both peripheral and central mechanical sensitization (*9*). Joint replacement surgery often alleviates this sensitization and pain, suggesting that the osteoarthritic joint drives these processes, even at late stages of disease (*9*). Interestingly, it has also been shown that individuals with lowered pain pressure thresholds to mechanical pressure at the knee are more likely to go on to develop persistent knee pain, suggesting that mechanical sensitization plays a key role in the early stages of osteoarthritis pain (*10*).

Nerve growth factor (NGF) has been suggested as a therapeutic target for osteoarthritis pain, and clinical trials with antibodies that neutralize NGF reported positive results in terms of pain relief (*11, 12*). Using rodent models of osteoarthritis, we and others have shown that a unique subset of nociceptors (‘silent nociceptors’) become responsive to mechanical stimuli (indicating peripheral sensitization) during the course of disease (*13-16*). The role of Piezo2 in this process is unclear, but a link between NGF and Piezo2 has been proposed as the mechanism of activation of ‘silent nociceptors’ (*17*). Specifically, transfection of silent nociceptors (*Chrna3*+ nociceptors) with *Piezo2* targeting siRNA was sufficient to block NGF-induced mechanical sensitization of these cells *in vitro*. Previous publications exploring the role of Piezo2 in pain have used mice in which the channel was deleted from all sensory neurons, and therefore behavioral measures of mechanical allodynia were difficult to assess, since these mice cannot move normally due to proprioception deficits (*2*). Here, we deleted *Piezo2* specifically from nociceptors by using Na_V_1.8-Cre mice, and used these mice to examine the role of Piezo2 in the development of mechanical sensitization in experimental osteoarthritis. Our results suggest that nociceptor expression of Piezo2 plays a key role in this process and cooperates with NGF-mediated signaling in producing nociceptor sensitization. Hence, Piezo2 may represent a novel therapeutic target for osteoarthritis pain.

## RESULTS

### Subhead 1: *Piezo2* can be specifically deleted from nociceptors in mice by using the marker *Scn10a*

While recent studies suggest that Piezo2 mediates mechanical allodynia in models of acute inflammation and nerve injury (*3, 4*), the role of Piezo2 in mediating mechanical pain associated with specific diseases in animals and humans has yet to be investigated. A major obstacle to such studies has been that the pan-sensory neuron *Piezo2* knock-out mice that have been used to date also show pronounced proprioceptive deficits, making it difficult to accurately assess evoked responses to mechanical stimuli (*2*). Indeed, *Piezo2* mRNA is expressed by a wide range of sensory neurons, as seen in single cell RNA sequencing (scRNAseq) analyses of mouse and human DRGs (*18, 19*). We used scRNAseq to confirm that *Piezo2* is expressed by many subsets of nociceptors (Fig. 1A, S1), and RNAscope analysis revealed a large degree of overlap between nociceptors, defined as *Scn10a*+ neurons (*Scn10a* is the gene that encodes Na_V_1.8) and *Piezo2*, with 38±2% of DRG neurons expressing both *Scn10a* and *Piezo2* (Fig. 1B,C). Therefore, we decided to create nociceptor-specific conditional *Piezo2* knock-out mice (*Piezo2*^CKO^), enabling us to study the functional implications of Piezo2 expression by these neurons in particular. We crossed Na_V_1.8-Cre mice (*20*) with Piezo2 loxp mice (*2*), in which exons 43-45 are flanked by loxp sequences. RNAscope confirmed the effectiveness of this strategy; co-expression of *Scn10a* and *Piezo2* was reduced to 6±1% of DRG neurons in homozygous *Piezo2*^CKO^ mice (Fig. 1B, 1C, S2, S3).

**Fig. 1.**
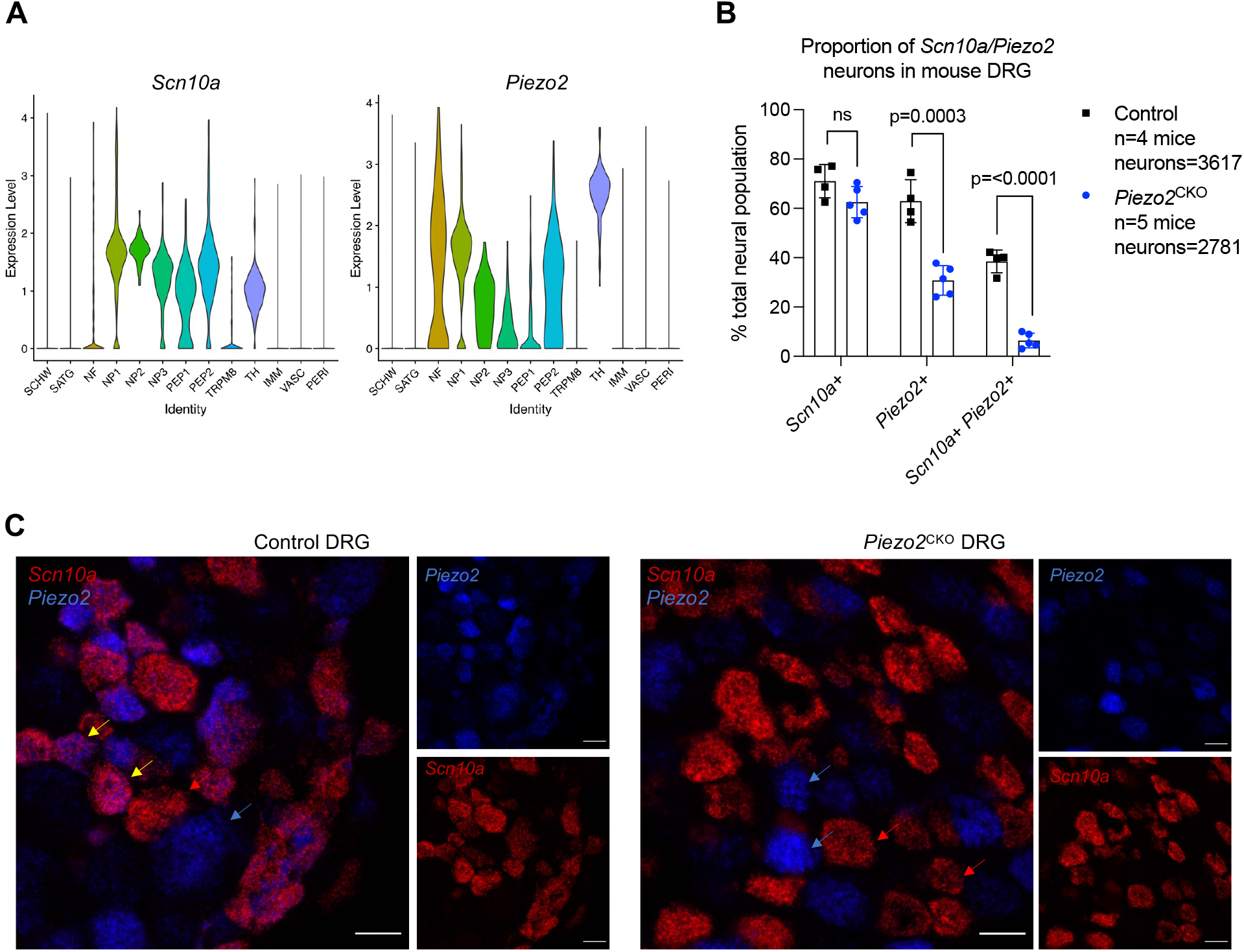
Piezo2 is deleted from nociceptors using the marker Na_V_1.8. A) ScRNA-seq of naïve L3-L5 DRG cells demonstrates that *Piezo2* is expressed by a subset of nociceptors (*Scn10a*+ neurons); B,C) *Piezo2* was deleted in nociceptors by using the marker Na_V_1.8 (gene *Scn10a*). RNAscope was used to quantify the number of cells expressing *Scn10a* and *Piezo2*. There is reduced co-expression of *Scn10a*+ / *Piezo2*+ in homozygous *Piezo2*^CKO^ mice (greater than 12 weeks of age). Unpaired t-test used to compare numbers of cells between strains. Mean±SD. C) Representative sections. Scale bar = 25 μm. Yellow arrow indicates example cells expressing *Scn10a* and *Piezo2*. Red arrow indicates example cells only expressing *Scn10a*. Blue arrow indicates example cells expressing only *Piezo2*.

Previous work has shown that *Piezo2* deletion alters C and Aδ nociceptor responses to mechanical stimuli *ex vivo* (*4*) and in select subclasses of nociceptors *in vivo* (*21, 22*). *Ex vivo*, electrophysiology using a skin-nerve preparation demonstrated that both C and Aδ fibers had reduced firing in response to mechanical stimuli in *Piezo2* knockout mice (*4*). *In vivo*, knockdown of Piezo2 in rat DRG via intrathecal injection of Piezo2 antisense oligodeoxynucleotides reduced Aδ fiber discharge frequency of bone afferents in response to pressure applied to the marrow cavity (*21*). Likewise, AAV mediated knock-out of Piezo2 in mice reduced responses by Aδ high-threshold mechanoreceptors to painful mechanical stimuli applied to the cheek as assessed by *in vivo* calcium imaging of the trigeminal ganglia (*22*). To confirm these observations, we used GCaMP6s expressing lines of the newly generated *Piezo2*^CKO^ mice (Na_V_1.8-Cre^+/-^;GCaMP6s loxp^fl/+^;Piezo2 loxp^fl/+^ mice) and performed *in vivo* calcium imaging of the L4 DRG to examine nociceptor responses to mechanical stimuli applied to the hindlimb of anesthetized mice (Fig. 2A). By imaging the L4 DRG we were able to observe the responses of a large population of sensory neurons that innervate the knee joint (*15, 23*). We applied two different levels of non-noxious mechanical force (30 g and 100 g) to the knee joints, and observed reduced numbers of neurons responding to these forces in naïve mice in which one copy of *Piezo2* was selectively deleted from nociceptors (representative images in Fig. 2B; Fig. 2C; Supplemental videos 1,2). In addition, the size of the responses quantified by peak area under the curve were reduced with the 100 g force in the *Piezo2*^CKO^ mice (Fig. 2D). Collectively, these data suggest that the *Piezo2*^CKO^ mice are suitable for studying the role of Piezo2 specifically expressed by nociceptors in disease models.

**Fig. 2.**
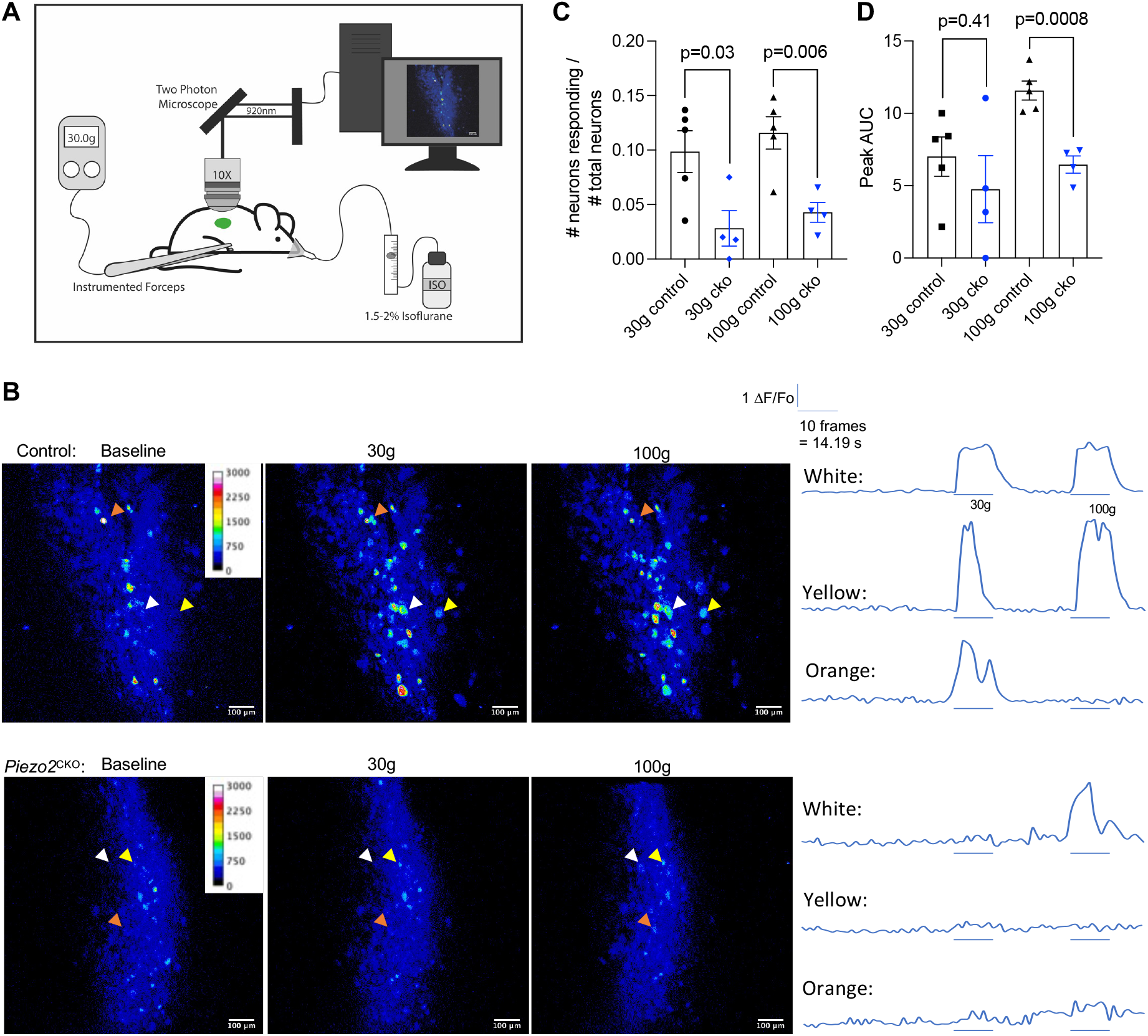
*In vivo* calcium imaging demonstrates a role for nociceptor expression of Piezo2 in mediating responses to non-noxious mechanical stimuli. (A) The L4 DRG is imaged by two photon microscopy in anesthetized mice while mechanical stimuli are applied to the knee joint using an instrumented forceps. (B) Representative DRG images and individual neuronal traces from control and *Piezo2*^CKO^ mice. Corresponds to supplemental video 1 and 2. (C) The number of Na_V_1.8+ neurons responding to each stimulus was quantified and compared between strains by unpaired t-test (each dot = one mouse; for each mouse >225 neurons imaged in total) (control, n=5 mice; *Piezo2*^CKO^, n=4 mice). (D) The peak area under the curve (AUC) of each neuron responding to each stimulus was quantified and averaged for each mouse. (strains compared by unpaired t-test; each dot = one mouse)

### Subhead 2: Deletion of *Piezo2* from nociceptors protects mice from osteoarthritis mechanical sensitization

Osteoarthritis pain is highly mechanical in nature, and sensitization to mechanical stimuli is a key feature of experimental osteoarthritis models as well as the human disease (*24*). Therefore, we decided to investigate if Piezo2 contributes to the development of mechanical sensitization associated with experimental osteoarthritis. To this end, we performed destabilization of the medial meniscus (DMM) surgery in Na_V_1.8-Cre mice and in homozygous *Piezo2*^CKO^ mice. We and others have previously characterized pain-related behaviors associated with mechanical input over the course of this slowly developing model of osteoarthritis. Mice show a reduced threshold for mechanically induced pain behaviors after DMM, including hyperalgesia at the injured knee and mechanical allodynia at the ipsilateral hind paw (*25-28*). Homozygous *Piezo2*^CKO^ were protected from both knee hyperalgesia and hind paw mechanical allodynia through 8 weeks after DMM (Fig. 3A,B), suggesting that Piezo2 expressing nociceptors are important for development of mechanical sensitization in this model, but that other mechanisms may drive mechanical allodynia by week 16. Joint damage was assessed 16 weeks after DMM, and showed that cartilage damage and osteophyte formation were comparable in *Piezo2*^CKO^ and control mice (Fig. S4). An independent experiment was performed using both heterozygous and homozygous *Piezo2*^CKO^ mice, demonstrating a similar pattern of protection against mechanical allodynia 4 and 8 weeks after DMM surgery (Fig. S5). Importantly, these mice did not exhibit signs of proprioceptive deficits due to *Piezo2* deletion (Fig. S5A) and had a normal body weight (Fig. S5B).

**Fig. 3.**
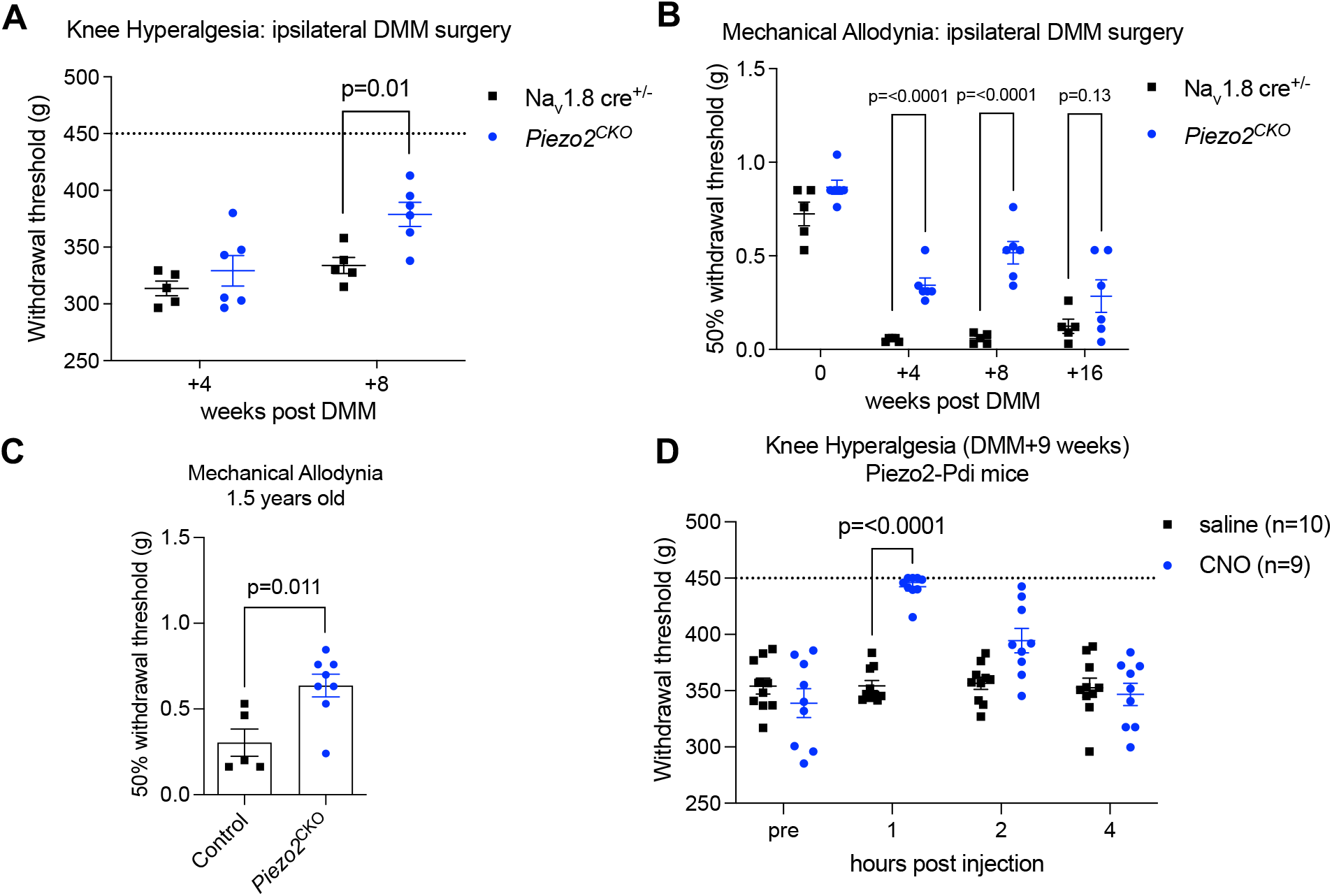
Piezo2 plays a role in mechanical sensitization in two mouse models of osteoarthritis. (A) Knee hyperalgesia was assessed in Na_V_1.8-Cre^+/-^ (n=5) and homozygous *Piezo2*^CKO^ (n=6) mice after DMM surgery. Dashed line indicates maximum of the assay: 450 g. Contralateral unoperated legs: 4 weeks (mean±SEM): Na_V_1.8-Cre^+/-^ (444±4) and *Piezo2*^CKO^ (440±10). 8 weeks: Na_V_1.8-Cre^+/-^ (443±4) and *Piezo2*^CKO^ (417±10). Two-way repeated measures ANOVA with Sidak post-test. (B) Hind paw mechanical allodynia was assessed in mice from part A. Two-way repeated measures ANOVA with Sidak post-test on log-transformed data. Independent experiment shown in Fig. S5. (C) Hind paw mechanical allodynia was assessed in unoperated control mice (littermate no Cre, n=5) or *Piezo2*^CKO^ (n=8) at 1.5 years of age. Unpaired t-test on log-transformed data. (D) Knee hyperalgesia in Piezo2-Pdi inhibitory DREADD mice 9 weeks after DMM surgery was assessed before, 1, 2, and 4 hours after intra-articular injection of saline (n=10 mice) or CNO (n=9 mice). Dashed line indicates maximum of the assay: 450 g. Two-way repeated measures ANOVA with Sidak post-test.

Age is a major risk factor for osteoarthritis (*8*), and male wild type as well as homozygous *Piezo2*^CKO^ mice develop age-associated cartilage damage and osteophyte formation in the knee joints by age 22 months (Fig. S6). Concordantly, at 18 months of age, these wild type mice had a reduced hind paw withdrawal threshold to mechanical stimuli (Fig. 3C) compared to young mice (time 0 in Fig. 3B = 10 weeks old). In contrast, *Piezo2*^CKO^ mice had a higher withdrawal threshold at 18 months of age compared to controls (Fig. 3C), once again indicating the role of this channel in mechanical sensitization in osteoarthritis.

Finally, to investigate whether targeting Piezo2 may represent a suitable strategy for osteoarthritis pain, we used chemogenetics to acutely silence *Piezo2*+ nerves within the knee joint (heterozygous Piezo2-Cre;Pdi mice). We found that intra-articular injection of clozapine-N-oxide (CNO) 9 weeks after DMM surgery transiently reversed knee hyperalgesia compared to saline (Fig. 3D), suggesting that local inhibition of *Piezo2*+ nociceptors represents a promising therapeutic target.

### Subhead 3: A subset of nociceptors co-express *Piezo2* and *Ntrk1* in mouse and human DRG

Mechanical sensitization has long been recognized as a major feature of osteoarthritis pain (*9*) and may be mediated by several inflammatory factors present in the osteoarthritis joint, including cytokines and growth factors that are synthesized and released as a result of ongoing tissue damage (*29*). In particular, the neurotrophin NGF, which is upregulated in osteoarthritic joint tissues (*30*), produces rapidly sensitizing effects in nociceptors through activation of its high-affinity receptor, tropomyosin receptor kinase A (TrkA) (*31*). A recent study observed that application of NGF to mouse DRG neurons in culture greatly increased their mechanosensitivity, and *Piezo2* siRNA inhibited this response (*17*). Additionally, mechanical hypersensitivity induced by local injection of NGF into bone marrow could be reduced by *Piezo2* knockdown in rats (*21*). Hence, a precise description of the Piezo2 and TrkA signaling axis in the joint is expected to increase our understanding of how pain is produced in osteoarthritis. RNAscope of the lumbar DRGs of adult healthy mice indicated that *Piezo2* and *Ntrk1* (gene that encodes TrkA) are co-expressed by 27±4% of Scn10a+ nociceptors (Fig. 4A-C, S7). Likewise, analysis by scRNA-seq also found co-expression of *Piezo2* and *Ntrk1* in 38% of *Scn10a*+ DRG neurons (Fig. S8A). In particular, the PEP2 cluster, which also includes the *Chrna3*+ silent nociceptor population, had the highest amount of co-expression (Fig. S8B).

**Fig. 4.**
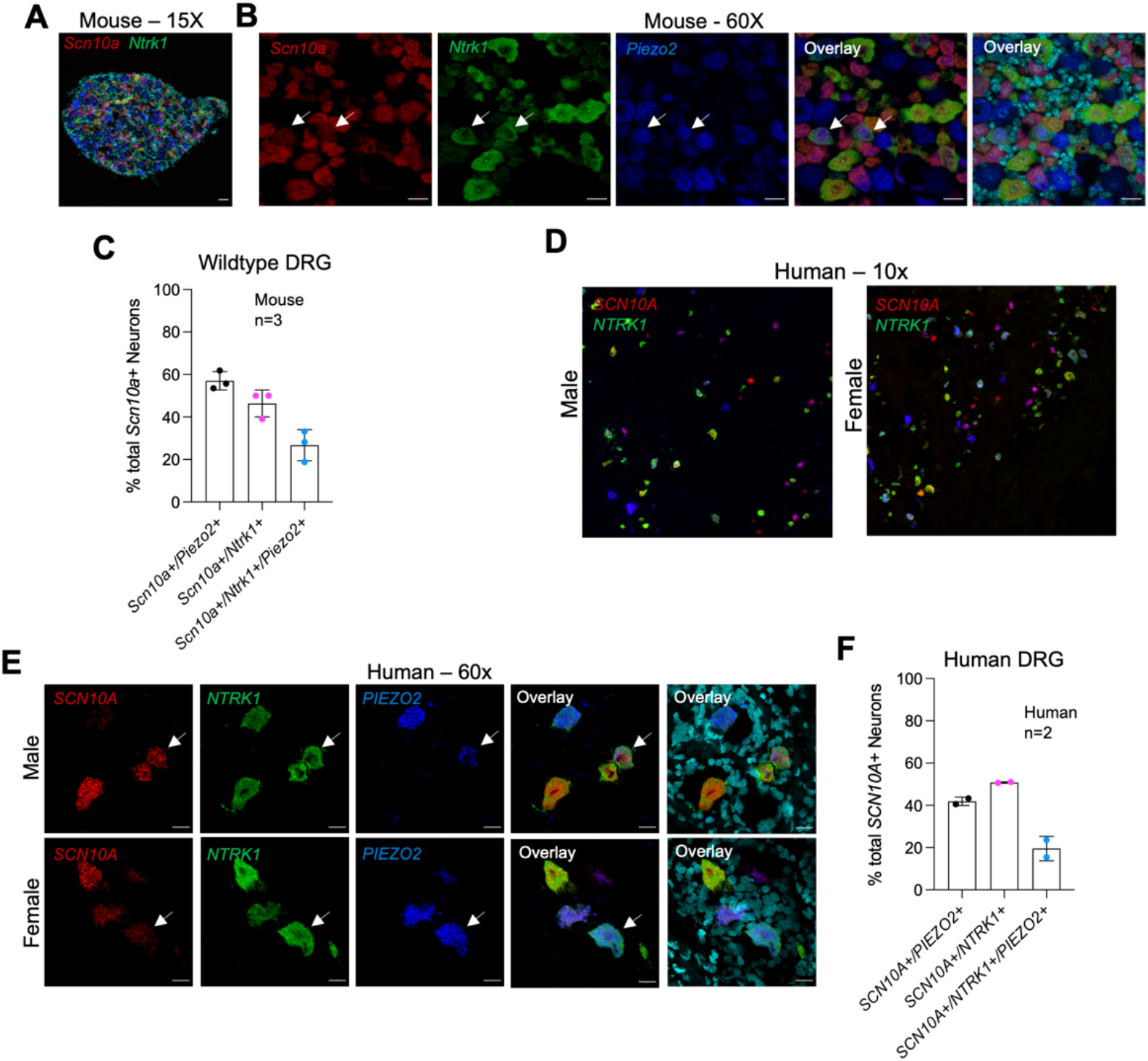
Subset of sensory neurons co-express *Scn10a, Piezo2*, and *Ntrk1* in both A-C) mouse and D-F) human DRG. (A,B,C) RNAscope used to quantify the number of cells expressing *Scn10a, Ntrk1* and *Piezo2* with DAPI in mouse DRG (n=3 mice) or (D,E,F) *SCN10A, NTRK1* and *PIEZO2* with DAPI in Human DRG (n=2 donors; one male and one female); Mean±SD. (A,B) Representative sections of mouse DRG. White arrows indicate example cells expressing *Scn10a, Ntrk1* and *Piezo2*. Scale bar = 50μm (A) and 25μm (B). (D,E) Representative sections of Human DRG. White arrows indicate example cells expressing *SCN10A, NTRK1* and *PIEZO2*. Scale bar = 50μm (D) and 25μm (E).

In human lumbar level DRG, a similar pattern of co-expression was observed by RNAscope. In DRGs collected *post mortem* from two donors (male: age 82; BMI 21.2 and female: age 86; BMI 27.6), examination of a total of 431 neurons showed that 42% of *SCN10A*+ neurons co-expressed *PIEZO2* (Fig. 4D-F, S9-11). In addition, *PIEZO2* and *NTRK1* were found to be co-expressed by 20% of *SCN10A*+ neurons (Fig. 4D-F, S9-11). This suggests that these co-expression patterns may be translationally relevant.

### Subhead 4: Deletion of Piezo2 blocks NGF-induced knee swelling and mechanical sensitization

Local injection of NGF into the hind paw or the knee joint has been shown to cause swelling as well as thermal and mechanical sensitization (*32-34*). The exact mechanisms by which NGF exerts these effects are not clearly understood, and NGF can potentially act through numerous mechanisms including enhancing release of inflammatory mediators, transactivating nociceptor ion channels, and longer-term effects on gene expression (*35*). NGF has also been shown to play an important role in mediating osteoarthritis pain (*30*). In order to examine how NGF directly impacts mechanical sensitization in the knee joint in concert with Piezo2 over an extended time period, and whether this involves activation of Piezo2, we modeled this interaction by injecting recombinant murine NGF bi-weekly (500 ng NGF in 5 μL) into the knee joint cavity of adult naïve wild-type or heterozygous *Piezo2*^CKO^ (Na_V_1.8-Cre^+/-^;GCaMP6s loxp^fl/+^;Piezo2 loxp^fl/+^) mice over the course of 8 weeks. This intra-articular treatment protocol produced knee swelling in wild-type naïve mice by day 7, and swelling was sustained through the end of the study on day 57 (Fig. 5A,B). In contrast, *Piezo2*^CKO^ mice were protected from NGF-induced knee swelling at all time points (Fig. 5A,B). Wild-type mice developed knee hyperalgesia by week 2, and this continued through week 8 (Fig. 5C,D). In contrast, while *Piezo2*^CKO^ mice also developed early knee hyperalgesia at the 2-week time point, these mice were protected from sustained knee hyperalgesia, consistent with their lack of knee swelling (Fig. 5C,D). Previously, mechanical and thermal sensitization induced by NGF has been linked to upregulation of neuropeptide expression in DRG neurons (*36*). To explore this possible link, we performed an additional experiment, in which we used qPCR to analyze the expression of *Calca* (the gene encoding CGRP) in the L3-L5 DRG after 3 intra-articular bi-weekly injections of vehicle or NGF. In wild-type mice, NGF injection resulted in increased DRG levels of *Calca* compared to vehicle, whereas *Piezo2*^CKO^ mice were protected from this effect (Fig. 5D). Together, these experiments support the idea that NGF induces its downstream effects in concert with Piezo2.

**Fig. 5.**
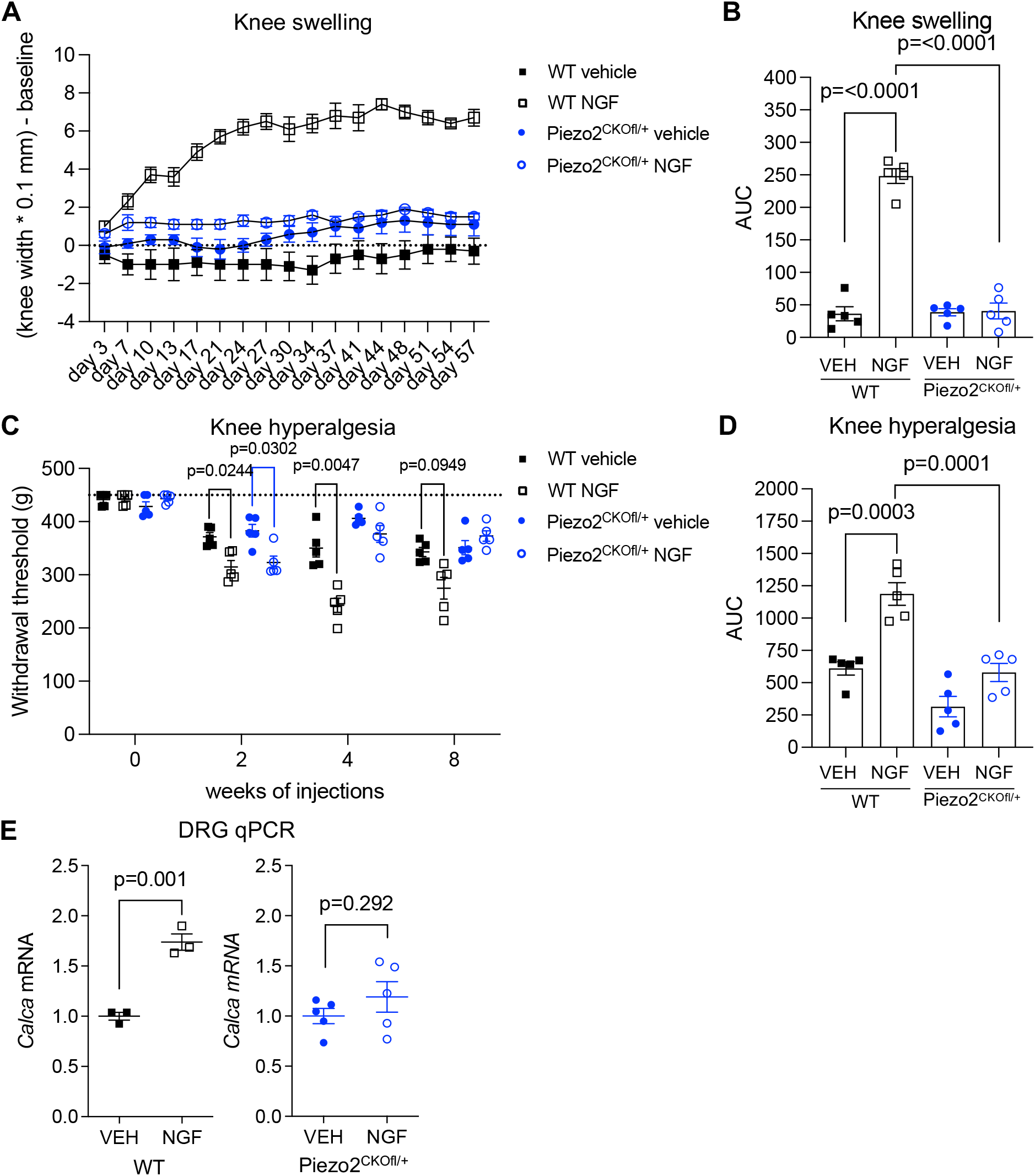
Piezo2 deletion protects from NGF induced knee swelling and mechanical sensitization. (A) Knee swelling was assessed in WT or heterozygous Piezo2^CKOfl/+^ mice given repeated intra-articular injections of recombinant murine NGF (i.a. 500 ng, 5 μL, 2x/week) or vehicle (5 μL, 2x/week) for 8 weeks; n=5 mice/group. Repeated measures two-way ANOVA with Sidak’s post-test: WT NGF vs vehicle, p<0.05 from day 7 onward. Piezo2^CKOfl/+^, no differences between NGF vs vehicle. (B) Area under the curve analysis over the time course was used to assess knee swelling. Two-way ANOVA with Sidak’s post-test. (*Piezo2*^CKO^ vehicle vs. *Piezo2*^CKO^ NGF: p>0.99). (C) Knee hyperalgesia was assessed in the same mice as part A. Dashed line indicates maximum of the assay: 450 g. Two-way repeated measures ANOVA with Sidak’s post-test. (D) Area under the curve analysis over the time course was used to assess knee hyperalgesia. Two-way ANOVA with Sidak’s post-test. (*Piezo2*^CKO^ vehicle vs. *Piezo2*^CKO^ NGF: p=0.12). (E) RNA was extracted from the L3-L5 DRGs of a separate cohort of mice after 3 injections of NGF or vehicle and qPCR was performed. Unpaired two-tailed t-test.

## DISCUSSION

Our observations support a key role for Piezo2 expressed by nociceptors in mediating mechanical sensitization associated with two mouse models of osteoarthritis, as well as with long-term mechanical sensitization induced by NGF administered locally into the knee. Consistent with this interaction, co-expression of *Piezo2* and *Ntrk1* was demonstrated in subsets of murine as well as human nociceptors. These results support previous work suggesting that Piezo2 is expressed by nociceptors (*4, 17, 21, 22, 37*). However, its primary function on nociceptors does not appear to be sensation of noxious mechanical forces under healthy conditions; rather Piezo2 appears to become sensitized in settings of inflammation and tissue damage. This supports previous work demonstrating that pan-sensory neuron deletion of *Piezo2* reduced mechanical pain following acute application of capsaicin and in a model of nerve injury (*4*). In addition, individuals with *PIEZO2* loss of function mutations had reduced mechanical allodynia following capsaicin application to the skin (*3*). Osteoarthritis is associated with mechanical sensitization and joint pain with movement. Joint damage products released as a result of ongoing tissue remodeling in osteoarthritis, including NGF, have been implicated in the development of mechanical sensitization (*24, 29*). Our findings indicate that Piezo2 is an essential component of this phenomenon.

Previous work has shown that *Piezo2* deletion alters C and Aδ nociceptor responses to mechanical stimuli *ex vivo* (*4*) and in select subclasses of nociceptors *in vivo* (*21, 22*). Here, by using *in vivo* calcium imaging of the DRG, we also observed that nociceptor responses to mechanical stimuli applied to the knee joint were decreased in anesthetized *Piezo2*^CKO^ mice. However, we found that nociceptor deletion of Piezo2 only impacts behavioral responses of mice after joint damage or inflammation has been initiated. This is consistent with other work suggesting that in the absence of inflammation, noxious mechanical stimuli are detected by an as yet unidentified channel/receptor (*22, 38, 39*).

Other work has shown that Piezo2 activity can be enhanced on short time scales by inflammatory molecules such as bradykinin (*40, 41*) and NGF (*17, 21*), but exactly which signaling pathways are involved remains unclear (*42-44*). On short time scales, NGF has been shown to trans-activate other channels such as TRPV1. However, *chronic* sensitization likely involves retrograde transport of NGF to the DRG, where it can promote changes in gene expression and/or membrane localization of channels and upregulation of neuropeptides. Previous work demonstrated that NGF-induced edema was in part generated through neuropeptide release by capsaicin-sensitive nociceptors (*32*). As in the current study, a mechanical component appeared to be involved in this effect since rats that were anesthetized had less edema than awake animals (*32*). We also found that *Piezo2* deletion inhibits NGF-induced DRG gene expression of *Calca*, the gene encoding the neuropeptide CGRP. Hence, our results suggest that chronic NGF-mediated sensitization of joint nociceptors, which is critical to osteoarthritic pain, is also dependent on Piezo2.

Our study has some limitations. Since we targeted nociceptors broadly by using Na_V_1.8-Cre mice, we did not functionally assess the impact of Piezo2 expressed by different subtypes of nociceptors. However, our transcriptional analyses help to inform the most likely subsets contributing to the observed phenotype. Both the ‘NP1’ and ‘PEP2’ subclasses of nociceptors had strong co-expression of *Scn10a* and *Piezo2*, while only the ‘PEP2’ type had strong co-expression with *Ntrk1* as well. This type of neuron also expresses the silent nociceptor marker, *Chrna3* (Fig. S8), supporting previous work that also implicated this type of sensory neuron with respect to NGF and Piezo2 signaling (*17*). In addition, recent comparative analyses of subsets of nociceptors in humans *vs*. mice has revealed that there are marked species differences in the exact subsets of nociceptors (*19, 45, 46*). While we show here using RNAscope that subsets of human DRG neurons do co-express *SCN10A, PIEZO2*, and *NTRK1*, more work must be done to understand the functional implications in humans.

To translate this work, development of specific therapeutic strategies to inhibit Piezo2 will be critical. A dietary fatty acid was recently shown to inhibit Piezo2 when activated by bradykinin *in vitro* (*41*), however the specificity of such approaches is still unclear. In osteoarthritis, intra-articular or topical therapies are attractive options for local delivery, which could avoid unwanted systemic effects of Piezo2 blockade associated with blocking proprioception (*2, 47*). It has been reported that intra-articular injection of the non-specific mechanosensitive channel inhibitor, Gsmtx4, was able to transiently inhibit joint pain in mice (*48*). Additionally, intra-articular delivery of Gi-DREADD via AAV was also shown to be effective in reducing inflammatory joint pain (*49*). Importantly the observation that inactivation of a single *Piezo2* allele in mice reduces pain associated with NGF has translational relevance since it was found that individuals with inactivating variants in *PIEZO2* in compound heterozygosity are protected from mechanical allodynia associated with application of capsaicin to the skin (*3*). This suggests that drugs that only partially block PIEZO2 may be effective and would thereby reduce problems associated with complete inhibition of PIEZO2 function.

## MATERIALS AND METHODS

## Study design

The objective of this study was to test the role of Piezo2 expressed by nociceptors in mechanical pain-related behaviors associated with experimental joint pain. To do this we used three different model systems: destabilization of the medial meniscus surgery, spontaneous osteoarthritis associated with aging, and long-term intra-articular injection of NGF, and we tested the development of pain-related behaviors in control mice and mice with *Piezo2* deletion in nociceptors. In addition, the expression pattern of *Piezo2* and the NGF receptor, *Ntrk1*, in nociceptors was examined in both mouse and human DRGs by RNAscope; additionally single cell RNAseq was used to examine mouse DRG expression patterns. For all experiments, the number of biological replicates and statistical test used are reported in the figure legends. For *in vivo* experiments, cages of mice were randomly assigned to treatment groups. Behaviors were assessed by an individual blinded to mouse genotype or experimental group. Sample sizes were chosen based on previous experience and publications in our laboratory. All animal experiments were approved by the Institutional Animal Care and Use Committees at Rush University Medical Center and Northwestern University.

## Animals

A total of 134 mice were used. Animals were housed with food and water *ad libitum* and kept on 12-hour light cycles. Na_V_1.8-Cre mice were obtained as a gift from Dr. John Wood (on C57BL/6 background) (*20*). tdTomato loxp (Jax # 007909) or GCaMP6s loxp (Jax # 028866) mice were ordered from Jackson labs and crossed with Na_V_1.8-Cre mice. Piezo2-Cre (Jax # 027719) and Piezo2 loxp (Jax # 027720) mice were ordered from Jackson labs and crossed with Na_V_1.8-Cre, Na_V_1.8-Cre;tdTomato or Na_V_1.8-Cre;GCaMP6s mice. hM4Di-loxp mice (inhibitory DREADD receptor) (termed Pdi mice) were obtained as a gift from Dr. Susan Dymecki (on C57BL/6 background) (*50*). Piezo2-Cre mice were crossed with Pdi mice to generate heterozygous Piezo2-Pdi mice, used for all chemogenetic experiments.

## Humans

Human DRGs came from participants in the Religious Orders Study (ROS) or Rush Memory and Aging Project (MAP) (*51, 52*). At enrollment, participants agreed to annual clinical evaluation and organ donation at death, including brain, spinal cord, nerve, and muscle. Both studies were approved by an Institutional Review Board of Rush University Medical Center. All participants signed an informed consent, Anatomic Gift Act, and a repository consent to allow their resources to be shared. The DRGs were removed postmortem and flash frozen as part of the spinal cord removal. One DRG was from a male (age 82; BMI 21.2) and one DRG was from a female (age 86; BMI 27.6). ROSMAP resources can be requested at https://www.radc.rush.edu.

## *In vivo* DRG calcium imaging

Naïve adult male mice, age 35-39 weeks (n=5 Na_V_1.8;GCaMP6s^fl/+^; n=4 Na_V_1.8;GCaMP6s^fl/+^;Piezo2^fl/+^) were used. All mice were deeply anesthetized using isoflurane (1.5-2% in O_2_), a laminectomy from vertebrae L2-L6 was performed, and the right-side L4 DRG was exposed (*15*). This DRG contains cell bodies of sensory neurons that innervate the mouse knee joint and hind paw (*53, 54*). Silicone elastomer (World Precision Instruments) was used to cover the exposed DRG and surrounding tissue to avoid drying (*55*). The mouse was positioned under a Prairie Systems Ultima In Vivo two photon microscope on a custom stage, using Narishige spinal clamps to slightly elevate the mouse in order to avoid motion artifacts associated with breathing. A Coherent Chameleon-Ultra2 Ti:Sapphire laser was tuned to 920 nm and GCaMP6 signal was collected by using a bandpass filter for the green channel (490 nm to 560 nm). Image acquisition was controlled using PrairieView software version 5.3. Images of the L4 DRG were acquired at 0.7 Hz, with a dwell time of 4 μs/pixel (pixel size 1.92 × 1.92 μm^2^), and a 10x air lens (Olympus UPLFLN U Plan Fluorite, 0.3 NA, 10 mm working distance). The scanned sample region was 981.36 × 981.36 μm^2^. Anesthesia was maintained using isoflurane (1.5-2%) during imaging. Mechanical force was applied to the mouse limb using a calibrated forceps device (*15*). For each mouse, baseline images were acquired in the absence of force (30 frames), followed by imaging while non-noxious forces of 30 g or 100 g were applied to the knee joint of the mouse. For each stimulus, baseline images were captured for 10 frames prior to the application of the stimulus, the stimulus was applied for 10 frames, and an additional 10 frames were captured after the stimulus was discontinued. Between each stimulus, the mouse was allowed to recover for at least 3 min in order to ensure that all previous neuronal responses had ceased and the fluorescence levels had returned to baseline. In addition, as a positive control, neuronal responses to a 200 g force applied to the ipsilateral hind paw were confirmed (*15*) prior to proceeding with the rest of the experiment. For each mouse, changes in [Ca^2+^]_i_ were quantified using a custom ImageJ macro to calculate the change in fluorescence in each frame t of a time series using the formula: ΔF/Fo, where Fo is the average intensity of the baseline period acquired in the absence of force (30 frames) (*15*). Sensory neuron responses to either 30 g or 100 g mechanical force applied to the knee joint were identified as cells having peak ΔF/Fo during the application period that was greater than 4 times the standard deviation of the baseline period (*55*). The total number of neurons imaged for each DRG was estimated by counting the number of neurons within a region of average density and extrapolating to the total imaged area (mean±SEM: Na_V_1.8;GCaMP6s = 360±42 neurons; Na_V_1.8;GCaMP6s;Piezo2^fl/+^ = 294±40 neurons). The percentage of responses to each stimulus was calculated using the formula: # responses / # total neurons imaged × 100. The peak area under the curve (AUC) of each neuron responding to each stimulus was quantified using GraphPad Prism and averaged for each mouse.

## Surgery

DMM surgery was performed in the right knee of 10-week old male mice (25 – 30 g), as previously described (*28, 56*), under isoflurane anesthesia. Briefly, after medial parapatellar arthrotomy, the anterior fat pad was dissected to expose the anterior medial meniscotibial ligament, which was severed. The knee was flushed with saline and the incision closed. Mice were not administered analgesia *post* surgery.

## Mechanical Allodynia

Control and *Piezo2*^CKO^ mice were tested for secondary mechanical allodynia of the ipsilateral hind paw using von Frey fibers and the up-down staircase method, as previously described (*28*). For the DMM experiment corresponding to Fig. 3B, S4: Na_V_1.8-Cre (n=5) and homozygous *Piezo2*^CKO^ (n=6) mice were tested. For the DMM experiment corresponding to Fig. S5: No Cre (n=7), *Piezo2*^CKOfl/+^ (n=7) and *Piezo2*^CKOfl/fl^ (n=10) mice were tested. Withdrawal thresholds were assessed before surgery and up to 16 weeks after DMM. For the aging experiment corresponding to Fig. 3C, S6: naïve no Cre (n=5) and naïve *Piezo2*^CKO^ mice (n=8) were tested at 18 months of age.

## Knee hyperalgesia

Knee hyperalgesia was measured using a Pressure Application Measurement (PAM) device (Ugo Basile, Varese, Italy), as previously described (*26*). Briefly, mice were restrained by hand and the hind paw was lightly pinned to make the correct flexion at a similar angle for each mouse. The PAM transducer was pressed against the medial side of the knee and pressure applied against the knee. PAM software guided the user to apply a constantly increasing force (30 g/s) up to a maximum of 450 g. If the mouse tried to withdraw its knee, the force at which this occurred was recorded. Two measurements were taken and recorded per knee and the withdrawal force data were averaged. Knee hyperalgesia was assessed 4 and 8 weeks after DMM surgery in Na_V_1.8-Cre^+/-^ (n=5) and *Piezo2*^CKO^ (n=6) mice by an experimenter blinded to the mouse strain.

## Inhibitory DREADD

Clozapine-N-oxide (CNO) (10 mg/kg in saline, i.a.) (Sigma-Aldrich, St Louis, MO, USA) or vehicle (phosphate buffered saline (PBS)) was administered to test the effect of neuronal inhibition on knee hyperalgesia in Piezo2-Pdi mice 9 weeks after DMM surgery (n=10 vehicle; n=9 CNO). Knee hyperalgesia was tested two hours prior to injection and at the indicated times *post* injection by a blinded observer.

## Histopathology of the knee

For the DMM experiment corresponding to Fig. 3A,B, S4: Na_V_1.8-Cre (n=5) and homozygous *Piezo2*^CKO^ (n=6) mice were taken down 18 weeks after DMM surgery and knees were collected and H&E stained. For the DMM experiment corresponding to Fig. S5: No Cre (n=7), *Piezo2*^CKOfl/-^ (n=7) and *Piezo2*^CKOfl/fl^ (n=10) mice were perfused 16 weeks post DMM, knees were collected and stained with toluidine blue. For the aging experiment corresponding to Fig. 3C, S6: 22-month old naïve no Cre (n=5) and naïve *Piezo2*^CKO^ mice (n=8) were taken down and knees were toluidine blue stained. Mid-joint frozen sections were collected and stained using standard methods as described (*57*). Knee sections were evaluated for cartilage degeneration, bone sclerosis and osteophyte width using modified Osteoarthritis Research Society International recommendations, as previously described (*58*).

## scRNAseq

L3-L5 DRGs were isolated and pooled from 10 C57BL/6 male naïve mice at 18 weeks of age and treated with 20 mg/mL collagenase for 15 min. This was followed by addition of digestion mix (1X TrypLE, 25 U/mL Papain, 1 mM DNase1 and 20 mg/mL collagenase/dispase) for 20 min. The cells were pipetted using a regular bore glass pipette for about ten times until no visible clumps remained. The cell suspension was then passed through a 40 *μ*m filter and the filtrate was centrifuged at 100 g for 5 min at 4°C. The cell pellet was resuspended in 0.5 mL aCSF (87 mM NaCl, 2.5 mM KCl, 1.25 mM NaH_2_PO_4_, 26 mM NaHC0_3_, 75 mM Sucrose, 20 mM Glucose, 1 mM CaCl_2_, 7 mM MgSO_4_) and 0.5 mL DRG media and gently layered on top of an Optiprep gradient (90 *μ*L Optiprep, 455 *μ*L aCSF and 455 *μ*L DRG media). The gradient with the cell suspension was centrifuged at 100 g for 10 min at 4°C and the process was repeated twice to remove any contaminating myelin debris. The pellet was then resuspended in 100 *μ*L of media along with 10 *μ*L DNase.

Cell number and viability were analyzed using a Nexcelom Cellometer Auto2000 with AOPI fluorescent staining method (92% viability). 9,400 naïve cells were loaded into the Chromium Controller (10X Genomics, PN-120223) on a Chromium Next GEM Chip G (10X Genomics, PN-1000120), and processed to generate single cell gel beads in the emulsion (GEM) according to the manufacturer’s protocol. The cDNA and library were generated using the Chromium Next GEM Single Cell 3’ Reagent Kits v3.1 (10X Genomics, PN-1000286) and Dual Index Kit TT Set A (10X Genomics, PN-1000215) according to the manufacturer’s manual. Quality control for the constructed library was performed by Agilent Bioanalyzer High Sensitivity DNA kit (Agilent Technologies, 5067-4626) and Qubit DNA HS assay kit for qualitative and quantitative analysis, respectively. The library was sequenced on an Illumina HiSeq 4000 sequencer with 2 × 50 paired-end kits using the following read length: 28 bp Read1 for cell barcode and UMI, and 90 bp Read2 for transcript. Sequencing reads were assembled and aligned against the mm10-2020-A mouse reference using Cell Ranger v6.0.0 (10x Genomics). Raw fastq files and the expression count matrix have been deposited on NCBI GEO (accession number GSE198485). The expression count matrix was analyzed using the Seurat v4.0.1 R package (*59*). In brief, cells were filtered (nFeature_RNA > 200 and percent.mt < 15), resulting in 8,755 cells for analysis. Data were log normalized, scaled, and the 2000 most variable features were identified. Clustering was performed using UMAP (dims 17, resolution = 1). Markers for each cluster were identified using the Wilcoxon rank-sum test integrated in Seurat. Clusters were manually annotated using previous DRG datasets as a guide (*18, 60*). Cells expressing combinations of *Scn10a, Piezo2*, and *Ntrk1* were identified as cells having expression levels of these genes > 0.1.

## DRG RNAscope

Mouse L3-L5 DRG were collected (n=4 no Cre controls; n=5 *Piezo2*^CKO^; n=3 wild-type mice), fixed in 4% paraformaldehyde (PFA), transferred to 30% sucrose solution for cryoprotection, embedded in OCT and cryo-sectioned onto slides. Human DRG were removed postmortem and flash frozen. DRGs were embedded in OCT and cryo-sectioned onto slides. During sectioning, both mouse and human slides were kept within the cryostat at -20°C before storage at -80°C.

RNA *in situ* hybridization (ISH) was performed using ACD Bio-Techne RNAscope Multiplex Fluorescent v2 Assay. For mouse DRG, manufacturer’s instructions were followed. For human DRGs, modifications were made to the protocol to preserve tissue integrity. Briefly summarized, slides were removed from -80°C and immediately submerged in 4% PFA on ice for 40 min. Dehydration was performed following 50%, 75% and two 100% ethanol washes for 5 min each. Hydrogen peroxide (3%) was applied for 10 min. Target retrieval was performed, reducing time in target retrieval buffer to 3 min followed by protease III incubation for 30 min. The remainder of the protocol was performed following manufacturer’s instructions. Probes were used at 1:50 dilution and Opal dyes from Akoya Biosciences were used at 1:100 dilution. A combination of opal dyes 520 (OP-001001), 570 (OP-001003) and 650 (OP-001005) were used. For mouse DRG, *Scn10a* (426011-C2), *Piezo2-E43-E45* (439971-C3) and *Ntrk1* (435791-C1) probes were used and ACD Bio-Techne DAPI. For human DRG, *SCN10A* (406291-C3), *PIEZO2* (449951-C2) and *NTRK1* (402631-C1) probes were used. For DAPI staining, Vectashield containing DAPI was used. ACD Bio-Techne positive and negative control probes were conducted for each species prior to start of work. Negative controls were included on every slide.

All imaging was performed using a Fluoview FV10i confocal microscope at 10x and 60x magnification. Multiple planes of focus were captured, but Z-stacks were not produced and instead the optimally focused image was chosen for processing and analysis. Laser intensity was used at ≤9.9% throughout. Images were processed and quantified using Fiji software. Only brightness and contrast tools were used to adjust images. For neural counts, all neurons were first counted manually using ‘Multi-point’ tool on phase contrast images. Counts of neurons singularly positive for each signal were then performed, followed by dual expressing and triple expressing neurons. For mouse DRGs, 3-4 sections per mouse were quantified and averaged. For human DRGs, 3 images per 2-3 sections of each DRG were quantified and averaged. The total number of neurons assessed is indicated in figures. For stitching of human DRG images in supplemental figure 9 and 10, Fiji ‘Stitching’ plugin was used (*61*).

## NGF intra-articular injections: Study 1

Recombinant murine NGF (500 ng R&D Systems cat. No.1156) or vehicle (0.1% BSA in PBS) in 5 μL was injected intra-articularly twice a week for 8 weeks into the right knee of naïve wild-type or heterozygous *Piezo2*^*CKO*^ (Na_V_1.8-Cre^+/-^;GCaMP6s loxp^fl/+^;Piezo2 loxp^fl/+^ mice) male mice 15-16 weeks old (n=5/group-total of 20 mice). Knee swelling was assessed once before injections started (baseline) and before each injection using a microcaliper. Knee hyperalgesia was assessed at baseline, and at 2, 4 and 8 weeks in a blinded way as described above. **Study 2:** Recombinant murine NGF (500 ng R&D Systems cat. No.1156) or vehicle (0.1% BSA in PBS) in 5 μL was injected intra-articularly twice a week for a total of 3 injections into the right knee of naïve wild-type or *Piezo2*^CKO^ (Na_V_1.8-Cre^+/-^;GCaMP6s loxp^fl/+^;Piezo2 loxp^fl/+^ mice) male mice 15-16 weeks old (n=3-5 mice/group). Ipsilateral L3-L5 DRG were collected from mice 24 hours after the last injection of NGF or vehicle. Gene expression of *Calca* was analyzed in the DRG cells using qRT-PCR. Briefly, total RNA was extracted using RNeasy mini kit (Qiagen, Hilden, Germany). Reverse transcription was performed using RT^2^ first strand kit (Qiagen). Quantification of mRNA was conducted using the SYBR Green qPCR master mix (Qiagen) and the RT^2^ primer assays (Qiagen) on a Bio-Rad CFX96 machine. The comparative ΔΔCT method was utilized for relative quantitation of gene levels of expression. *Gapdh* was used as an internal control for normalization of target gene expression.

## Statistics

For RNAscope, counts were compared between genotypes by unpaired two-tailed t-test. For *in vivo* calcium imaging, the number of sensory neuron responses to a mechanical stimulus were compared between strains of mice by unpaired two-tailed t-test. For mechanical allodynia data, paw withdrawal thresholds were log-transformed prior to further analyses (*62*). For mechanical allodynia and knee hyperalgesia time courses, a repeated measures two-way ANOVA with Sidak post-test was used to compare Na_V_1.8-Cre and *Piezo2*^CKO^ responses at each time point. For mechanical allodynia in aged mice, genotypes were compared by unpaired two-tailed t-test. For knee hyperalgesia analgesic time courses, a repeated measures two-way ANOVA with Sidak post-test was used to compare mice treated with vehicle to mice treated with CNO at each time point. For knee histopathology, continuous data were analyzed by unpaired two-tailed t-test or one-way ANOVA with Sidak post test and non-continuous data were analyzed by Mann-Whitney test or Kruska-Wallis followed by Dunn’s post test. All analyses were carried out using GraphPad Prism version 9.3.1 for Mac (GraphPad Software, San Diego, CA). Results are presented as mean ± SEM unless otherwise noted in the figure legend. Sample size, p value, and statistical tests are indicated in the figure legends.

## Supporting information

SuppMaterial

## Acknowledgements

This work was supported by the Northwestern University NUSeq Core Facility. The 10x Genomics Chromium System employed for the scRNA-seq is made available with an NIH S10 Grant to NUSeq (1S10OD025120). We thank the study participants and staff of the Rush Alzheimer’s Disease Center. ROSMAP resources can be requested at https://www.radc.rush.edu.

## Author contributions

Conceptualization: AMO, MJW, RJM, AM, REM

Methodology: AMO, MJW, SI, LW, DR, RJM, AM, REM

Software: REM

Investigation: AMO, MJW, SI, JL, LW, DR, REM

Resources: DAB

Visualization: AMO, MJW, REM

Funding acquisition: RJM, AM, REM

Project administration: RJM, AM, REM

Supervision: RJM, AM, REM

Writing – original draft: AMO, MJW, RJM, AM, REM

Writing – review & editing: AMO, MJW, SI, JL, LW, DR, DAB, RJM, AM, REM

## Data and materials availability

Single cell RNAseq data have been deposited in NCBI GEO (accession number GSE198485).

## Supplementary Materials

Fig. S1. Additional single cell RNAseq information associated with Figure 1.

Fig. S2. RNAscope pictures from Figure 1.

Fig. S3. Additional RNAscope sections associated with Figure 1.

Fig. S4. Histology associated with DMM experiment in Figure 3.

Fig. S5. An independent experiment showing that Piezo2 plays a role in mechanical sensitization in the DMM mouse model of osteoarthritis.

Fig. S6. Histology associated with aging experiment in Figure 3. Fig. S7. Mouse DRG – additional sections from Figure 4.

Fig. S8. Single cell RNAseq data associated with Figure 4.

Fig. S9. Stitched image of male human DRG – corresponding to Figure 4.

Fig. S10. Stitched image of female human DRG – corresponding to Figure 4.

Fig. S11. Additional 60x images of human DRGs from Figure 4 to demonstrate different co-expression combinations of *SCN10A, NTRK1*, and *PIEZO2*.

Movie S1. Video showing baseline, 30 g and 100 g responses of the control mouse DRG associated with Fig. 2B.

Movie S2. Video showing baseline, 30 g and 100 g responses of the *Piezo2*^CKO^ mouse DRG associated with Fig. 2B.

