## Supplementary material for "Piezo2 expressing nociceptors mediate mechanical sensitization in experimental osteoarthritis": SuppMaterial

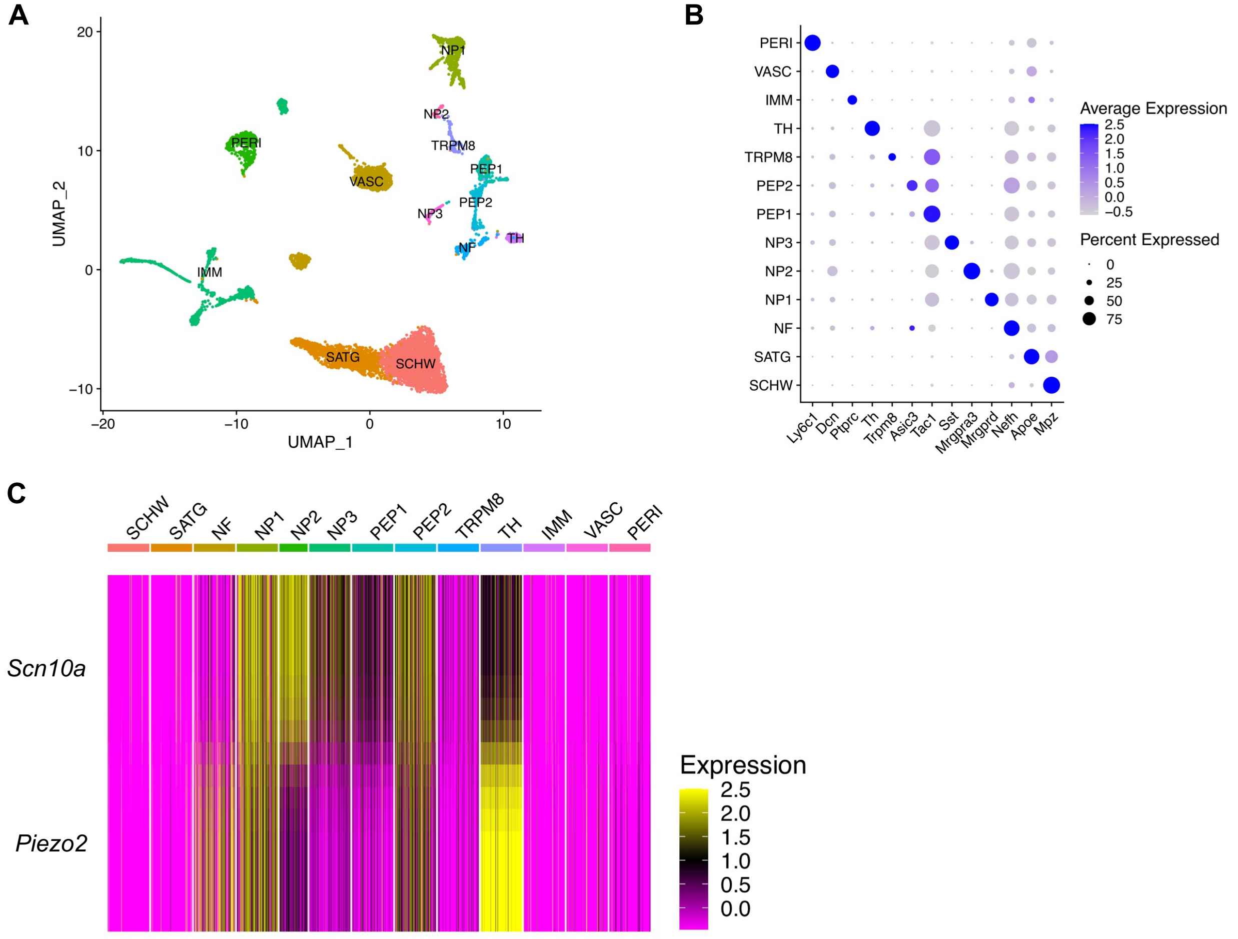


**D**

| Gene | Cluster | p_val | avg_log2FC | pct.1 | pct.2 | p_val_adj |
|---|---|---|---|---|---|---|
| *Piezo2* | NF | 2.58E-51 | 1.6244646 | 0.635 | 0.199 | 5.22E-47 |
| *Piezo2* | NP1 | 0 | 1.55685918 | 0.898 | 0.152 | 0 |
| *Piezo2* | PEP2 | 2.94E-88 | 0.75857994 | 0.829 | 0.193 | 5.94E-84 |
| *Piezo2* | TH | 2.55E-165 | 2.75989056 | 1 | 0.195 | 5.15E-161 |

**Fig. S1. Additional single cell RNAseq information associated with Figure 1.** (A) UMAP plot of 8,755 L3-L5 dorsal root ganglia (DRG) cells isolated from 18-week old male naïve C57BL/6 mice. Cells were clustered and labeled based on previously published datasets (*18, 60*). 'SCHW' = Schwann cells; 'SATG' = satellite glia; 'IMM' = immune cells; 'PERI' = pericytes; 'VASC' = vascular cells; remaining clusters = neurons following annotation strategy as in (*18, 62*). (B) Example marker genes used to identify each cluster. (C) Heatmap displaying expression of *Scn10a* and *Piezo2* on a cell-by-cell basis. (D) FindAllMarkers command in Seurat used to identify clusters with upregulated *Piezo2* expression. NF = *Scn10a*- neurons; NP1, PEP2, TH = *Scn10a*+ neurons.

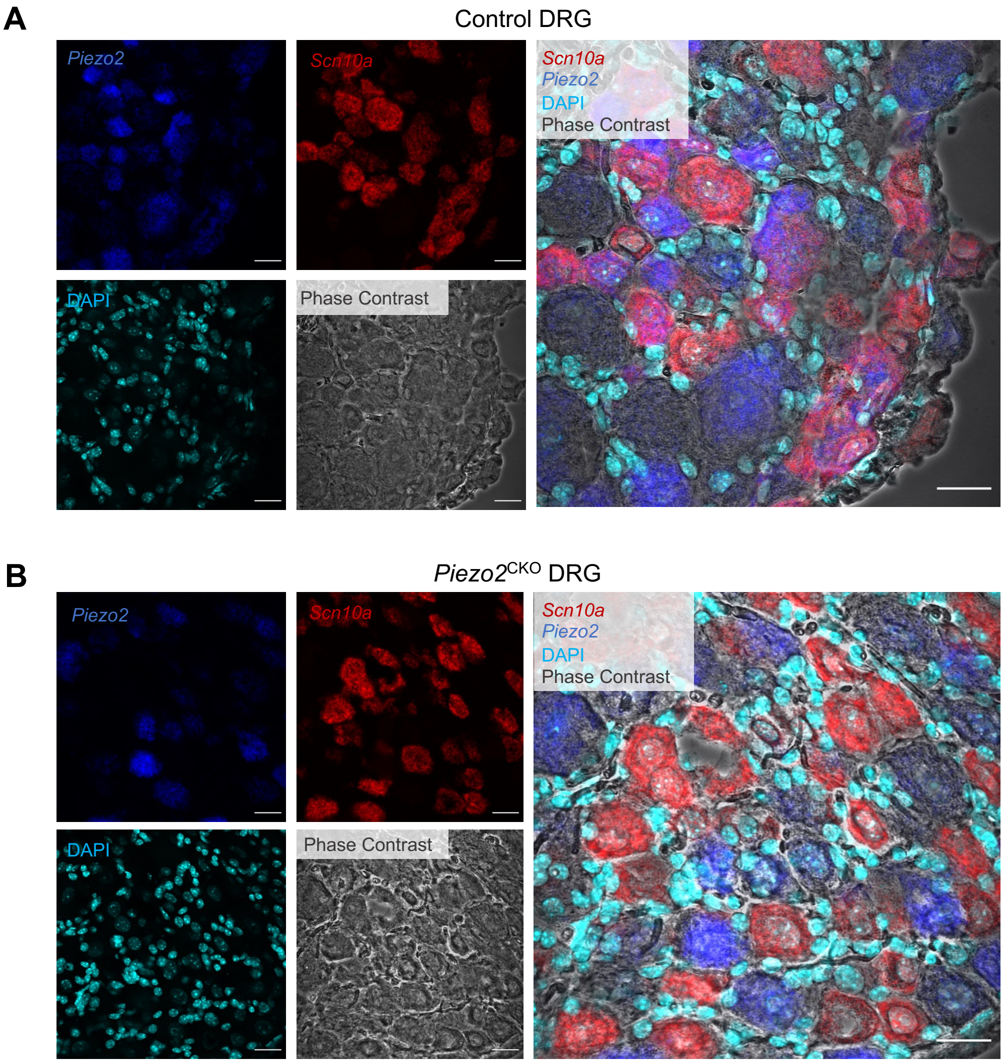


**Fig. S2. RNAscope pictures from Figure 1** in (A) control or (B) *Piezo2*$^{CKO}$ DRG with phase contrast and DAPI channels shown. Scale bar = 25 µm.

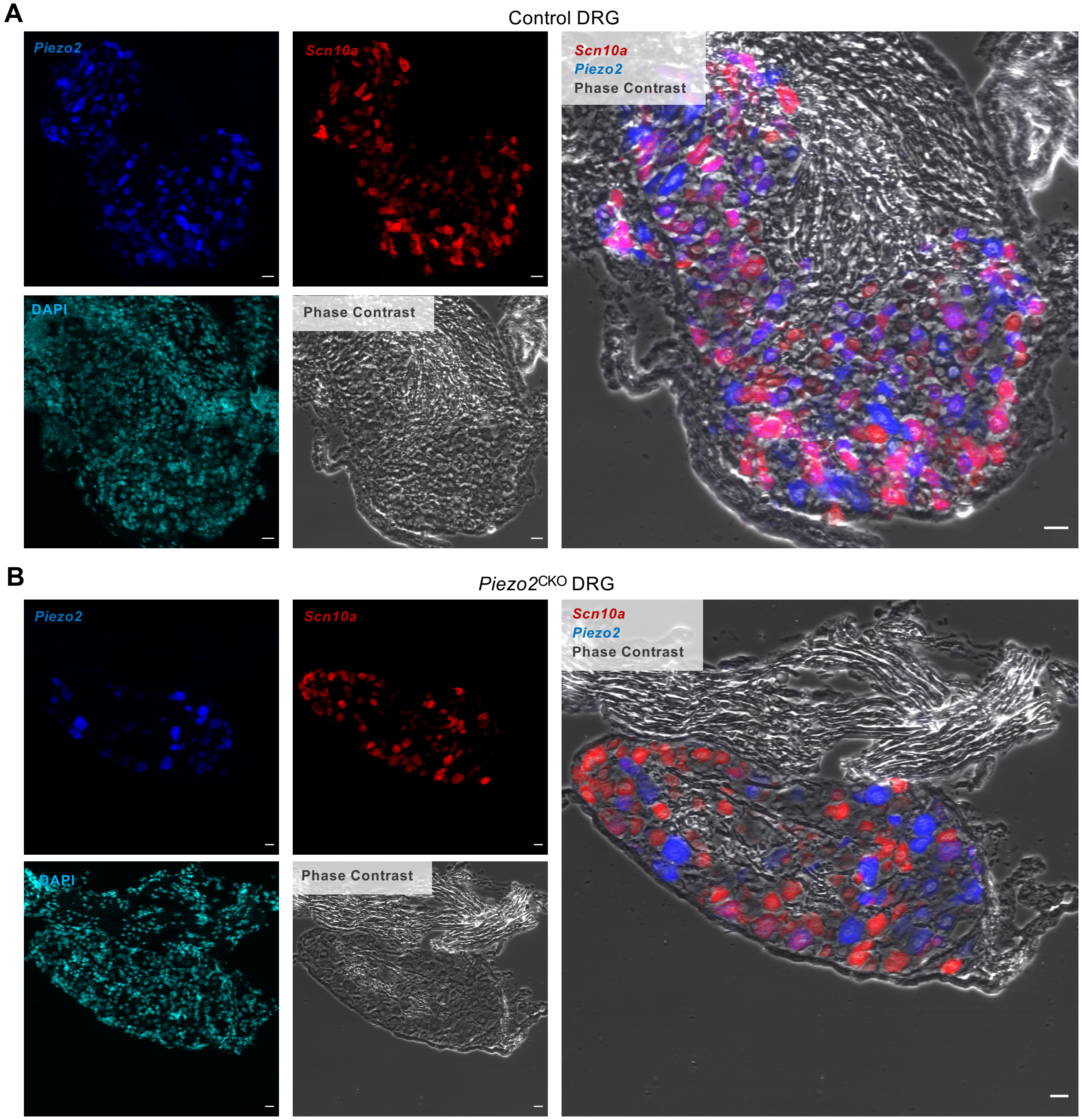


**Fig. S3. Additional RNAscope sections associated with Figure 1** in (A) control or (B) *Piezo2*$^{CKO}$ DRG with phase contrast and DAPI channels shown. Scale bar = 25 µm.

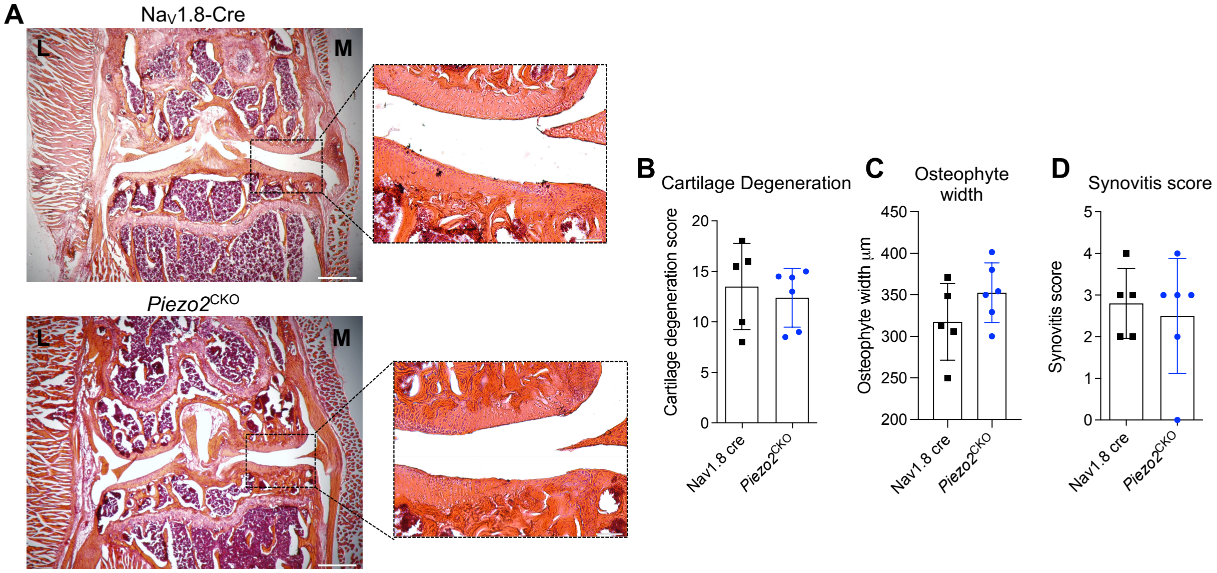


**Fig. S4. Histology associated with DMM experiment in Figure 3.** (A) Representative H&E histological images of the right knee joint 18 weeks after DMM surgery of $Na_V1.8$-Cre (n=5) and *Piezo2*$^{CKO}$ (n=6) mice from the experiment corresponding to Fig. 3A,B. L=lateral, M=medial side of the knee joint. Scale bar for whole joint image on left = 500 µm. Scale bar for inset = 100 µm. (B-D) represents modified OARSI scoring for cartilage degradation (unpaired two-tailed t-test, p=0.62), osteophyte width (unpaired two-tailed t-test, p=0.19) and synovitis score (Mann-Whitney, p=0.99) in the medial compartment, respectively. Mean±SD.

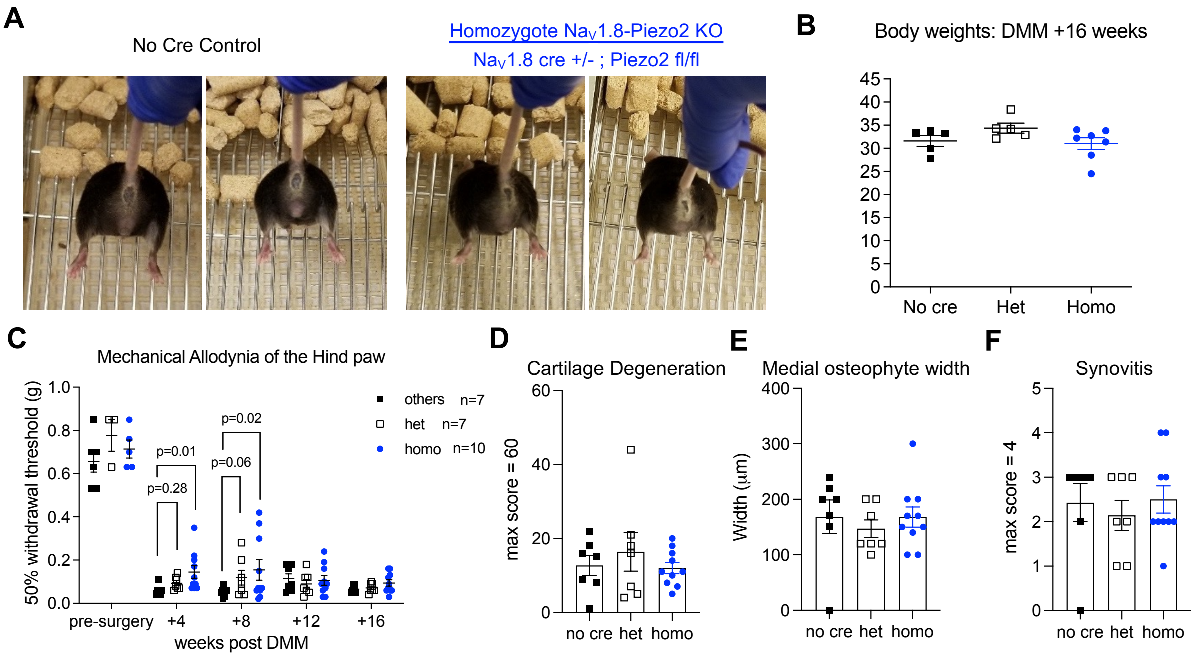


**Fig. S5. An independent experiment showing that Piezo2 plays a role in mechanical sensitization in the DMM mouse model of osteoarthritis.** (A) No effect on proprioception was detected with deletion of *Piezo2* from nociceptors using $Na_V1.8$ cre. (B) No change in body weight was detected with deletion of *Piezo2* from nociceptors (littermate no cre controls, n=5 mice), (heterozygous *Piezo2*$^{CKOfl/+}$, n=5 mice), (homozygous *Piezo2*$^{CKOfl/fl}$, n=7 mice). (C) Hind paw mechanical allodynia was assessed on log-transformed data by two-way ANOVA with Sidak post-test (littermate no cre controls 'others', n=7), (*Piezo2*$^{CKOfl/+}$, n=7), (*Piezo2*$^{CKOfl/fl}$, n=10). (D) Cartilage degeneration was assessed by one-way ANOVA with Sidak post-test (littermate no cre controls, n=7), (*Piezo2*$^{CKOfl/+}$, n=7), (*Piezo2*$^{CKOfl/fl}$, n=10). No cre vs. het: p=0.69; no cre vs. homo: p=0.98. (E) Osteophyte width was assessed by one-way ANOVA with Sidak post-test (littermate no cre controls, n=7), (*Piezo2*$^{CKOfl/+}$, n=7), (*Piezo2*$^{CKOfl/fl}$, n=10). No cre vs. het: p=0.77; no cre vs. homo: p=0.99. (F) Synovitis was assessed by Kruskal-Wallis test with Dunn's post-test (littermate no cre controls, n=7), (*Piezo2*$^{CKOfl/+}$, n=7), (*Piezo2*$^{CKOfl/fl}$, n=10). No cre vs. het: p=0.77; no cre vs. homo: p>0.99.

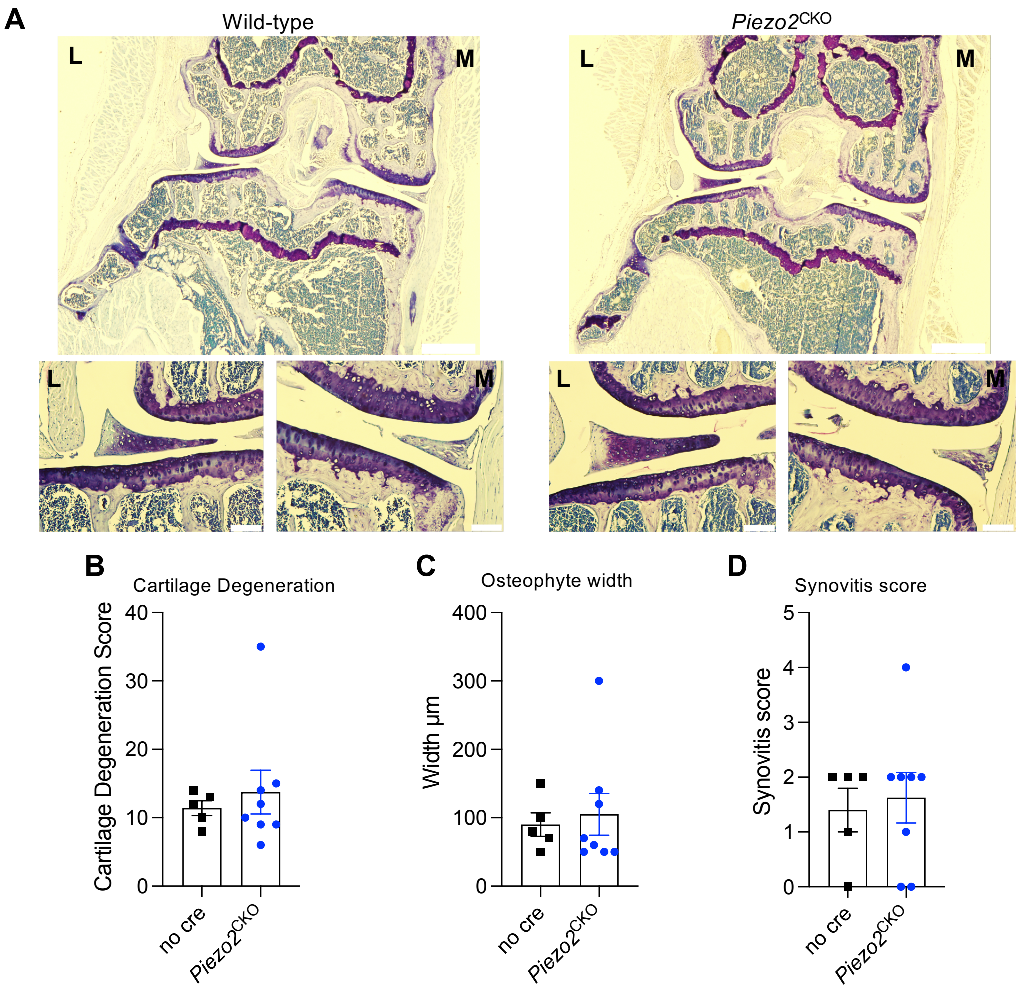


**Fig. S6. Histology associated with aging experiment in Figure 3.** (A) Representative toluidine blue histological images of the right knee joint of 22-month-old naïve mouse of littermate no cre controls (n=5) and *Piezo2*$^{CKO}$ (n=8) mice from the experiment corresponding to Fig. 3C. Scale bar for whole joint image on top = 500 µm. Scale bar for inset = 100 µm. (B-D) represents modified OARSI scoring for cartilage degradation, osteophytes width and synovitis score, respectively. (B) Unpaired t-test: no cre vs. *Piezo2*$^{CKO}$: p=0.59; (C) Unpaired t-test: no cre vs. *Piezo2*$^{CKO}$ : p=0.72; (D) Mann-Whitney test: no cre vs. *Piezo2*$^{CKO}$ : p=0.92. M = medial; L = lateral. Mean±SEM.

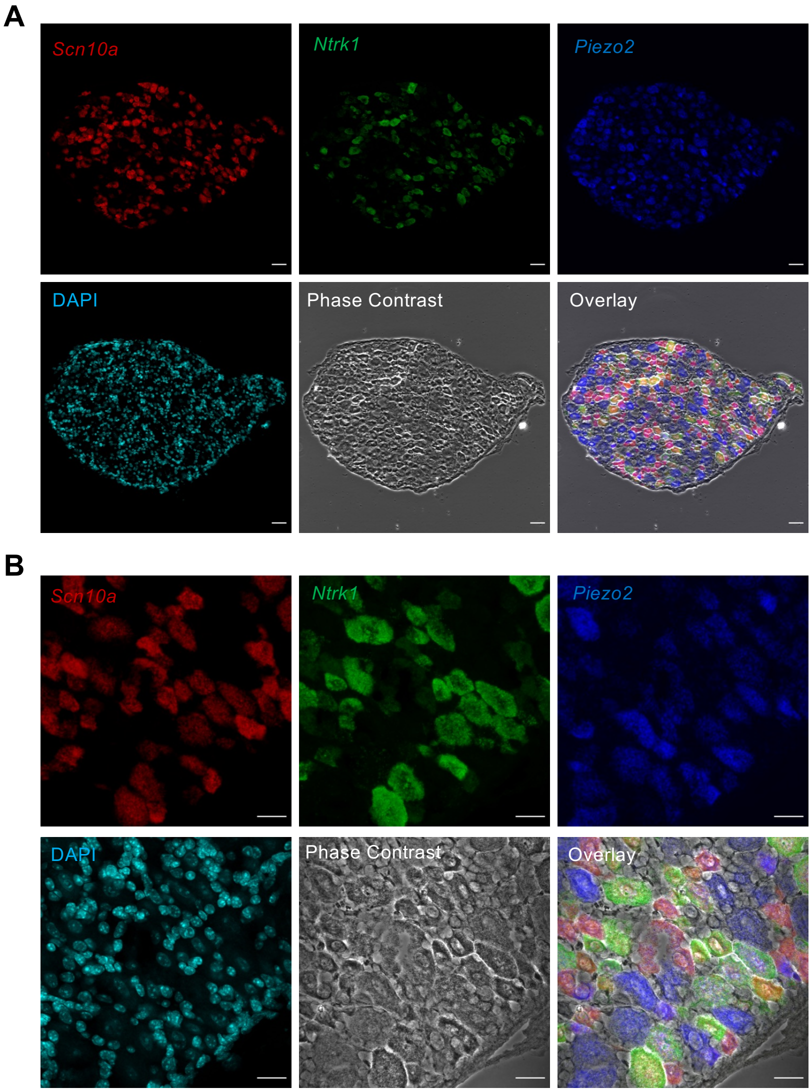


**Fig. S7. Mouse DRG – additional sections from Figure 4** at (A) 15X and (B) 60X magnification with phase contrast channel shown. Scale bar = 50 µm (A) and 25 µm (B).

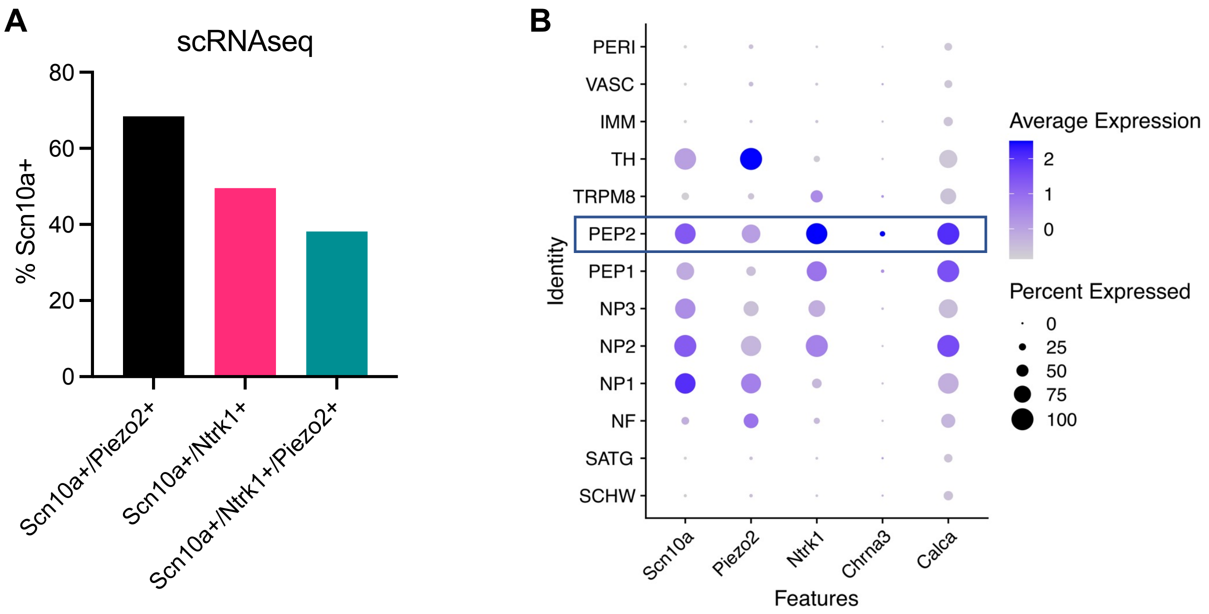


**Fig. S8. Single cell RNAseq data associated with Figure 4.** (A) Single cell RNAseq of mouse DRG also demonstrates co-expression of *Scn10a*, *Piezo2* and *Ntrk1*. (B) PEP2 cluster demonstrates strongest co-expression of *Scn10a*, *Piezo2* and *Ntrk1*; this cluster also expresses *Chrna3*, a marker for silent nociceptors, and *Calca* (gene that encodes CGRP).

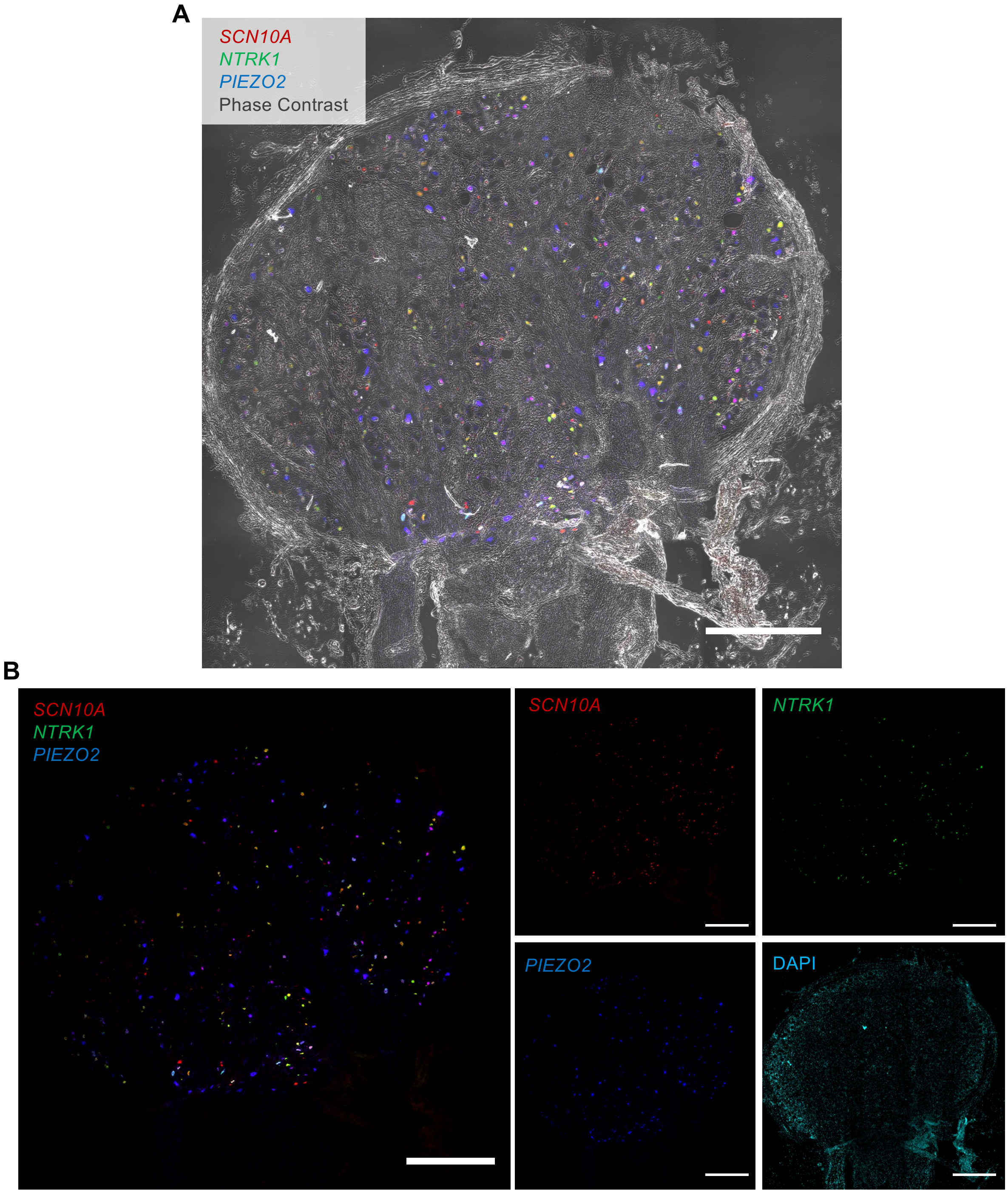


**Fig. S9. Stitched image of male human DRG – corresponding to Figure 4** showing (A) phase contrast and (B) individual channels of *SCN10A, PIEZO2, NTRK1* and DAPI. Scale bar = 1 mm.

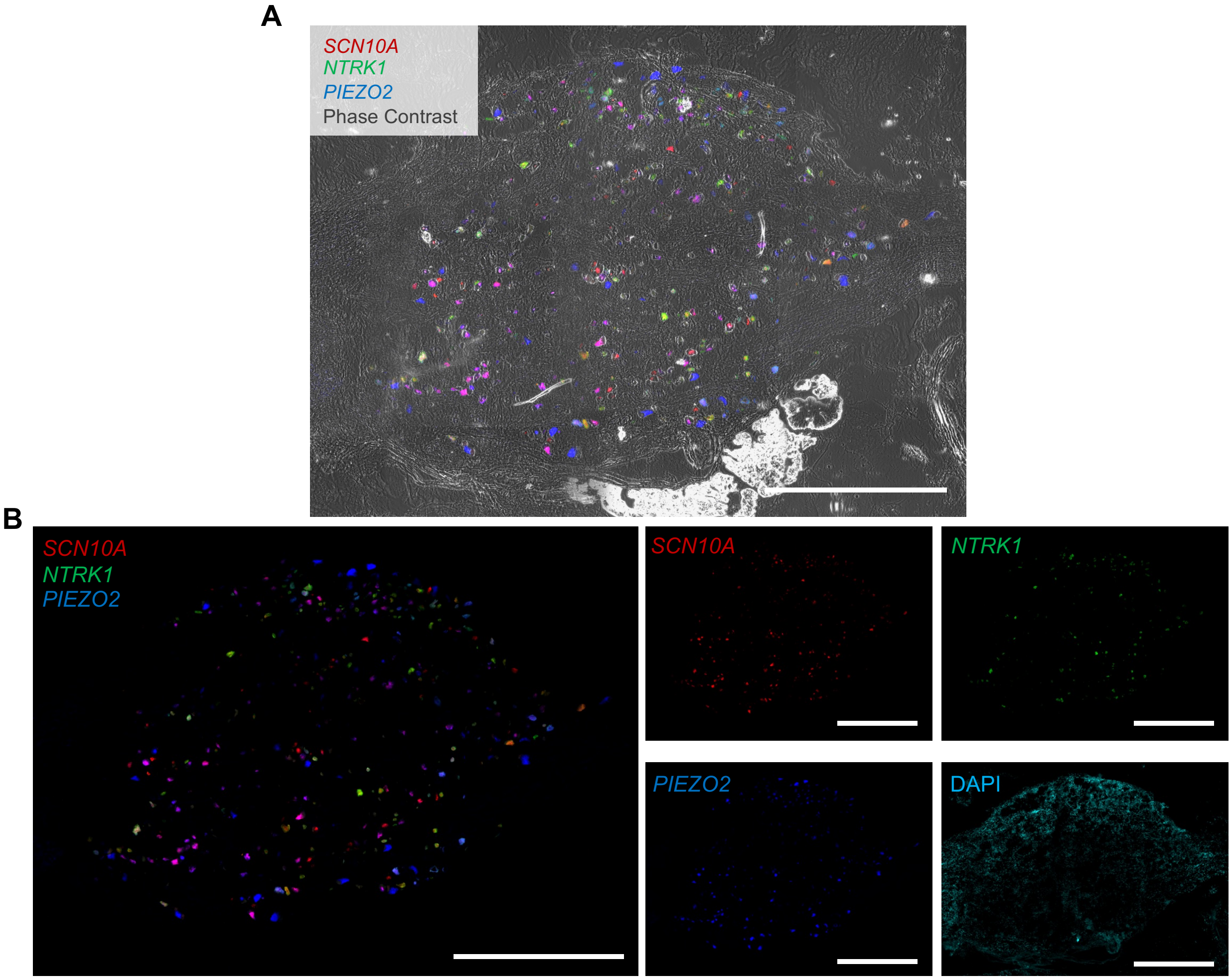


**Fig. S10. Stitched image of female human DRG – corresponding to Figure 4** showing (A) phase contrast and (B) individual channels of *SCN10A, PIEZO2, NTRK1* and DAPI. Scale bar = 1 mm.

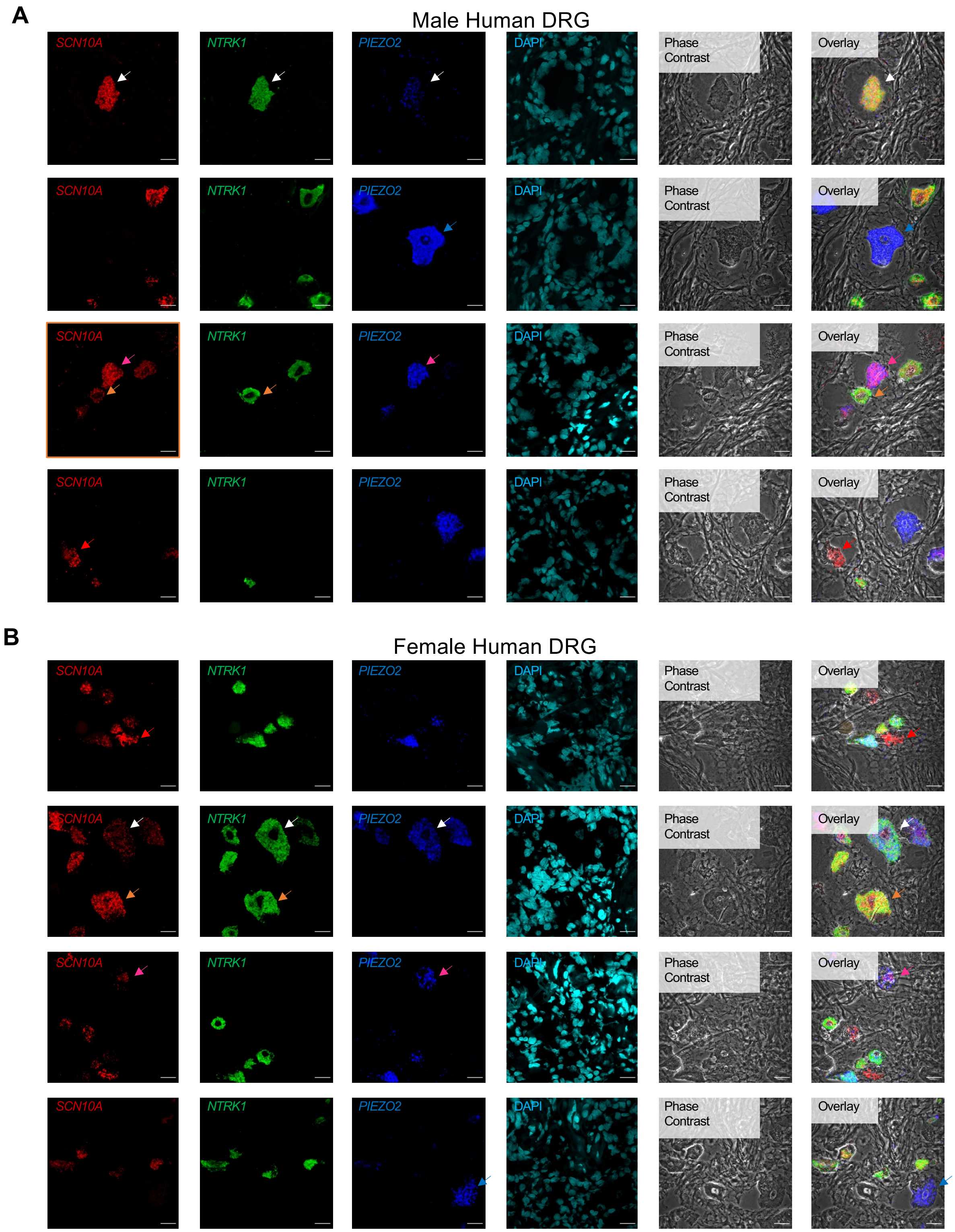


**Fig. S11. Additional 60x images of human DRGs from Figure 4 to demonstrate different co-expression combinations of *SCN10A, NTRK1*, and *PIEZO2*** in (A) male and (B) female human DRGs. White arrows indicate cells expressing *SCN10A, NTRK1*, and *PIEZO2.* Pink arrows indicate cells expressing *SCN10A* and *PIEZO2.* Orange arrows indicate cells expressing *SCN10A* and *NTRK1*. Red arrows indicate cells expressing only *SCN10A*. Blue arrows indicate cells expressing only *PIEZO2.* Scale bar = 25μm.
